# Male linked genomic regions determine sex in dioecious *Amaranthus palmeri*

**DOI:** 10.1101/2020.05.25.113597

**Authors:** Cátia José Neves, Maor Matzrafi, Meik Thiele, Anne Lorant, Mohsen B. Mesgaran, Markus G. Stetter

**Affiliations:** Institute for Plant Sciences, University of Cologne, Cologne, Germany; Dept. of Plant Sciences, University of California, Davis, CA, USA; Department of Plant Pathology and Weed Research, Agricultural Research Organization, Newe Ya’ar Research Center, Israel

**Keywords:** Amaranthus, dioecy, sex chromosome, invasive weed

## Abstract

Dioecy, the separation of reproductive organs on different individuals, has evolved repeatedly in different plant families. Several evolutionary paths to dioecy have been suggested, but the mechanisms behind sex determination is not well understood. The diploid dioecious *Amaranthus palmeri* represents a well suited model system to study sex determination in plants. *A. palmeri* is one of the most troublesome weeds in the US, has successfully colonized other regions in the world and has evolved resistance to several herbicide classes. Despite the agricultural importance of the species, the genetic control and evolutionary state of dioecy in *A. palmeri* is currently unknown. Early cytogenetic experiments did not identify heteromorphic chromosomes. Here, we used whole genome sequencing of male and female pools from two independent populations to elucidate the genetic control of dioecy in *A. palmeri*. Read alignment to a close monoecious relative and allele frequency comparisons between male and female pools did not reveal significant sex linked genes. Consequently, we employed an alignment free k-mer comparison which enabled us to identify a large number of male specific k-mers. We assembled male specific contigs comprising a total of almost 2 Mb sequence, proposing a XY sex determination system in the species. Based on our findings we suggest an intermediate evolutionary state of dioecy in *A. palmeri*. Our findings give insight into the evolution of sex chromosomes in plants and may help to develop sustainable strategies for weed management.

## Introduction

**T**he separation of sexes is observed in many animal and plant species, but while it is the norm in animals it is very rare in plants (Bachtrog *et al.* 2014). In most angiosperms, individuals have both functional sex organs and are therefore either hermaphrodite, i.e. having flowers comprising the reproductive organs of both sexes, or they are monoecious, where each plant carries distinct male and female flowers. However, in a small number of plant species (≈6% of known angiosperms (Renner 2014)) the female and male functions are harbored on separate individuals, i.e. they are dioecious (reviewed in Charlesworth 2016). Although rare, this breeding system has evolved independently in numerous taxonomic groups with 50% of plant families having at least one dioecious species (Renner 2014). A potential evolutionary advantage of dioecy over cosexuality is the avoidance of inbreeding and its resulting inbreeding depression. In addition, it allows the efficient allocation of resources to specialized functions of the respective sex (Charlesworth 2016).

Sex determination in dioecious plants can be environmentally or genetically controlled (Korpelainen 1998). The genetic control of sex determination in plants ranges from few individual genes (Spigler *et al.* 2008; Akagi *et al.* 2014) to non-recombining heteromorphic sex chromosomes (Sakamoto *et al.* 1998; Sousa *et al.* 2013). Most heteromorphic systems are XY male and XX female, but female specific sex chromosomes (ZW) also exist, e.g. in *Silene* and *Salix* (Slancarova *et al.* 2013; Pucholt *et al*. 2015). Although the genetic control of dioecy has been studied in several species, the determination of sex and the evolutionary forces driving sexual dimorphism are not well understood.

Functionally, the evolution of dioecy from a cosexual ancestral state requires at least two mutations, one creating males and one creating females (Westergaard 1958; Charlesworth and Charlesworth 1978). The simultaneous emergence of both mutations is highly unlikely. Hence, full dioecy has probably evolved via intermediate states, in which some individuals are female, while the others are still cosexual (Charlesworth 2016). The second mutation, which transforms the cosexual state into males by dominantly suppressing female functions, is likely occur in close linkage to the first mutation. This reduces recombination in the genomic region, as recombinant individuals would be sterile (Charlesworth 2016). In the most common case, where males are the heterozygous sex, lack of recombination between X and Y chromosomes creates a male-specific region of the Y chromosome, which can further result in gene degeneration (Ming *et al.* 2011).

As the non-recombining region spreads, further degeneration is accompanied with an accumulation of transposable elements (TEs), duplications and other mutations, leading to an increase in DNA content of the Y chromosome (Ming *et al.* 2011). In plant species with heteromorphic sex chromosomes, the Y chromosome is therefore usually bigger than their X counterpart (Charlesworth 2013; Westergaard 1958; Sakamoto *et al.* 1998; Sousa *et al.* 2013). Progressing degeneration can finally lead to massive gene loss in the Y chromosome and eventually a shrinkage of the sex chromosome. This last step is common in mammals (Graves 2006), and has been observed in plants (Abraham and Mathew 1962; Segawa *et al.* 1971), but appears to be less frequent (Bachtrog 2013). The above described signature of progressive sex chromosome evolution allows the identification of the evolutionary stage of sex determination from genomic data.

The genus *Amaranthus* comprises over 40 species (Sauer 1957), including crops grown for grains and vegetables (Joshi *et al.* 2018), as well as many invasive weeds of agricultural and natural systems (e.g. Rowland *et al.* 1999; Bensch *et al.* 2003). Most *Amaranthus* species are monoecious, while a few species exhibit a dioecious breeding system. In early classifications, dioecious *Amaranthus* species were grouped into a single subgenus (*Acnida*) based on their reproductive system (Mosyakin and Robertson 1996). Genome-wide phylogenetic work, however, indicated that these species cluster with monoecious species in distinct clades (Stetter and Schmid 2017). Hence, dioecy probably evolved multiple times independently within the *Amaranthus* genus.

Few weedy *Amaranthus* species are receiving increasing attention for their rapid evolution and spread of herbicide resistance (Kreiner *et al.* 2019; Molin *et al.* 2020). The dioecious *A. palmeri* L. is native to North America (Sauer 1957) and is one of the most devastating weeds in US agriculture (Webster and Nichols 2012; Riar *et al.* 2013). *A. palmeri* has evolved resistance to six difference classes of herbicides (reviewed in Ward *et al.* 2013, and www.weedscience.com) and is capable of producing copious amounts of seeds (up to 1 million per plant). Understanding the reproductive system and the sex determination of the species could help to develop agronomic strategies to decrease weed populations and mitigate herbicide resistances. Cytological analysis has shown that the chromosomes of male and female *A. palmeri* plants do not show heteromorphism (Grant 1959), suggesting an early stage of the evolution of dioecy in the species. Recent work using molecular markers suggested a genetic basis for sex determination in *A. palmeri*. However, due to low data quality and non-reproducible analyses, no inference of the evolutionary state of sex determination could be made (Montgomery *et al.* 2019). In addition, environmental factors can alter the expressed sex of an individual (Mesgaran *et al.* 2019).

In this study, we evaluated the evolutionary state of dioecy in *A. palmeri* to understand the genetic control and evolution of the separation of reproductive organs in plants. We show that dioecy in *A. palmeri* is controlled by a male specific genome region suggesting an XY system. We use high depth whole genome sequencing to distinguish between a sex gene system and male specific regions. Our high depth whole genome sequencing data reveals a 2 Mb sex specific region that could not be identified through allele frequency differences when aligned to a hermaphrodite relative, suggesting the ongoing evolution of a non-recombining male specific Y chromosome in *A. palmeri*.

## Materials and Methods

### Plant material

*A. palmeri* seeds of two independent populations from California and Kansas were used. Seeds of *A. palmeri* from California (CA) and Kansas (KS) were cordially provided by Dr. Anil Shrestha (California State University, Fresno, California) and Dr. Dallas E. Peterson (Kansas State University, Manhattan, Kansas), respectively. We grew seeds from the two populations under controlled well-watered conditions in the greenhouse in Davis, Caliofonia, USA, in the summer of 2018. About 10 seeds were planted in plastic pot (2.37 L), filled with a soil mix (1:1 sand/peat) plus a controlled-release. Plants were grown in a greenhouse with a temperature of 32/22°C (day/night) and a day length of 16 hours provided through supplementary lighting. Seedling were thinned randomly several times to obtain one plant per pot by their four-leaf stage. Plants were irrigated through four emitters inserted into the potting medium to deliver 65 mL of water min-1 for two minutes and twice per day (7:00 am and 2:00 pm). We collected equal amounts of leaf tissue (3 disks of 5 mm diameter) of each plant once the plants were flowering and the sex could be visually determined. We pooled the leaf samples of 35 male and 32 female plants from California and 25 male and 35 female plants from Kansas for DNA extraction.

### Sample preparation and sequencing

We extracted DNA from lyophilized and homogenized tissue using the DNeasy Plant Mini kit (Qiagen Sciences Inc, USA) and quantified DNA concentration on Qubit (Thermo Fisher Scientific, USA). Whole genome sequencing libraries were prepared by the UC Davis Genome Center for DNA Technologies & Expression Analysis core facility using the TruSeq DNA library kit (Illumina, USA). After quality control, the samples were sequenced on an Illumina NovaSeq. We assessed the data quality using fastqc (https://www.bioinformatics.babraham.ac.uk/projects/fastqc/) and estimated the sequence coverage based on the estimated genome size of 423 Mb (Stetter and Schmid 2017).

### Allele frequency comparisons

We aligned the raw reads to the monoecious *A. hypochondiracus* L. reference genome v 2.1 (Lightfoot *et al.* 2017) using BWA-MEM (Li 2013) and removed duplicates with picard (http://broadinstitute.github.io/pi Then, we called SNP using GATK (McKenna *et al.* 2010) with the filter expression `QD < 2.0 || FS > 60.0 || MQ < 40.0 || MQRankSum < −12.5 || ReadPosRankSum < −8.0` and further filtered to keep only biallelic SNPs with a maximum of 30% missing values at a site using VCFtools (Danecek *et al.* 2011). We created a variant table using `VariantsToTable` function of GATK to test for allele frequency differences between female and male pools using the QTLseqR package (Mansfeld and Grumet 2018).

### Reference free k-mer analysis

For the reference free comparison between male and female individuals we counted unique k-mers in the male and female pool from California. We first trimmed the reads based on their quality using trimmomatic (Bolger *et al.* 2014) with the parameters: `SLIDINGWINDOW:4:15 MINLEN:35 LEADING:5 TRAILING:5`, before counting the occurrences 35-mers in the quality trimmed sequencing reads from female and male pools separately keeping only k-mers with a frequency between 15 and 2000 using Jellyfish v 2.3.0 (Marçais and Kingsford 2011). We combined the male and female counts to identify sex specific k-mers using custom scripts. We further, counted sex specific k-mers identified in the Californian population in the pools of the Kansas population using `jellyfish query`. To further generate the most robust set of k-mers, we filtered sex specific k-mers requiring a k-mer to have a count ≥ 50 but ≤ 500 within the focal Kansas (e.g., female pool for female specific) and ≤ 20 in the Kansas opposite pool (e.g., male pool for female specific)

### Male specific read recovery

We selected male specific k-mers that had a count of minimum 50 in the male pool and 0 (represents ≤ 15 or ≥ 2,000, due to previous filtering) in the female pool. For all four sequenced pools, we extracted full size reads containing the k-mer sequences and their respective read pair from the trimmed read data using a custom bash script.

### Estimation of the size of male specific regions

To estimate the size of the male specific region we assembled the recovered male specific reads from the male California pool with platanus-allee v 2.0.2 (Kajitani *et al.* 2019) using default parameters. We counted the total length of assembled base pairs as sex specific region.

### Extraction of male k-mer containing reads from A. tuberculatus

We extracted the filtered set of male specific k-mers from published *A. tuberculatus* samples with known sex expression (Kreiner *et al.* 2019), proceeding as described above in “male specific read recovery”. To normalize the numbers we devided the number of extracted reads by the total number of sequenced reads.

## Results

### High coverage WGS of male and female pools

We applied high depth whole genome sequencing to four pools of *A. palmeri* from two independent wild populations, separated by sex (i.e. two populations x two sexes). The pools consisted of combined leaf samples of 35 male and 32 female individuals from a single population collected from California, as well as 25 male and 35 female individuals from another population collected from Kansas. The per pool sequence coverage ranged from 228X to 331X (322,106,487 to 467,707,106 reads per pool), corresponding to a mean coverage of 7.5X to 9.5X per sample (Table S1).

### No allele frequency differences between male and female pools

Given the lack of cytological evidence for heteromorphic sex chromosomes in *A. palmeri* (Grant 1959), we tested whether sex determination in *A. palmeri* could be controlled by a specific small and still recombining region, located on an autosome. To assess this possibility, we called biallelic SNPs and compared allele frequencies between the male and female pools of the two populations and tested for frequency differences. We aligned raw reads to the reference genome of the hermaphrodite *A. hypochondiracus* (Lightfoot *et al.* 2017), as no reference genome of *A. palmeri* is available, yet. Overall, reads mapped to the reference genome with 90.54% to 92.55% uniquely mapped reads, indicating a high similarity between the genomes of the two species (Table S1). Yet, a large number of read pairs showed non-proper paring, indicating structural differences between the genomes.

Sex determining loci in the genome are expected to lead to significant allele frequency differences between male and female pools. We used the G’ method implemented in QTLseqR to scan the first 16 Scaffolds of the genome for significant differences in allele frequencies. Using over 12 million biallelic sites (3.2 million after filtering), distributed across the genome, we found no significant allele frequency differences between male and female pools in either of the two populations (Figs. 1 and S1). Allele frequency differences between sexes were very low across the whole aligned genome sequence.

**Figure 1.**
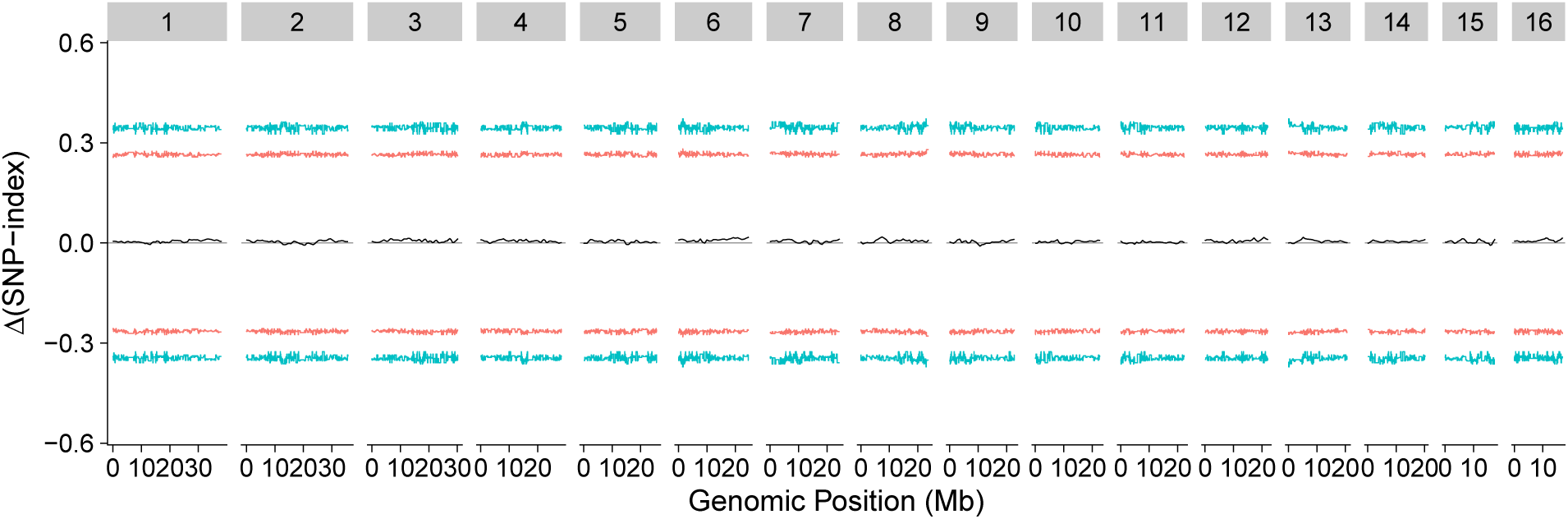
Differences in allele frequencies between male and female pools of the California population along the genome, relative to the *A. hypochondiracus* reference. Red and blue lines represent 95% and 99% confidence intervals for frequency outliers.

### Large number of male specific k-mers

Comparing allele frequency differences requires the successful alignment of reads to the reference genome. However, sequencing reads from diverged sex specific sequences are unlikely to align to the hermaphrodite reference genome as they accumulate large numbers of mutations (Ming *et al*. 2011; Charlesworth 2013). To test whether a larger, non-recombining, region could be responsible for sex determination in *A. palmeri*, we compared alignment free k-mer counts of male and female pools. We recovered all 35-mers (in the following referred to as k-mers) from the sequencing reads and counted the occurrences of these k-mers for each pool. Low frequency (≤ 15) and high frequency (≥2000) were discarded from the analysis as they potentially represent sequencing errors and repeat regions. We identified over 1.3 billion (1,329,452,869) male and over 1.2 billion (1,235,093,537) female k-mers that passed out filters and were used for further analysis. We compared k-mer counts of the male and female pools from the California population to find k-mers that strongly deviated from the expected equal frequency in male and female pools (Fig.2). As expected, most k-mers were present in comparable numbers within males and females (Fig.2). We found 1,634,859 k-mers that were not present in females (15 ≥n ≥2000), but had a count between 50 and 500 in the male pool. We used the Kansas population to obtain the most robust set of specific k-mers. That is, we counted and identified sex specific k-mers for the California population in the Kansas population. We further filtered male (or female) specific k-mers for a minimum count of 50 and a maximum of 500 in the Kansas pool (or female). To be considered sex specific we further required a count below 20 k-mers in the female (or male) Kansas pool. After these stringent filters, we found 158,693 robust male specific k-mers, which represent 9.7 % of all male specific k-mers (tab. S1). We also found a low number of female specific k-mers (269,157) in the Californian population. After correction with the Kansas population, we find 2,572 female specific k-mers (1.0 %, tab S1). These include few k-mers, that are present over 2000 times in the male pools and therefore not counted in males.

**Figure 2.**
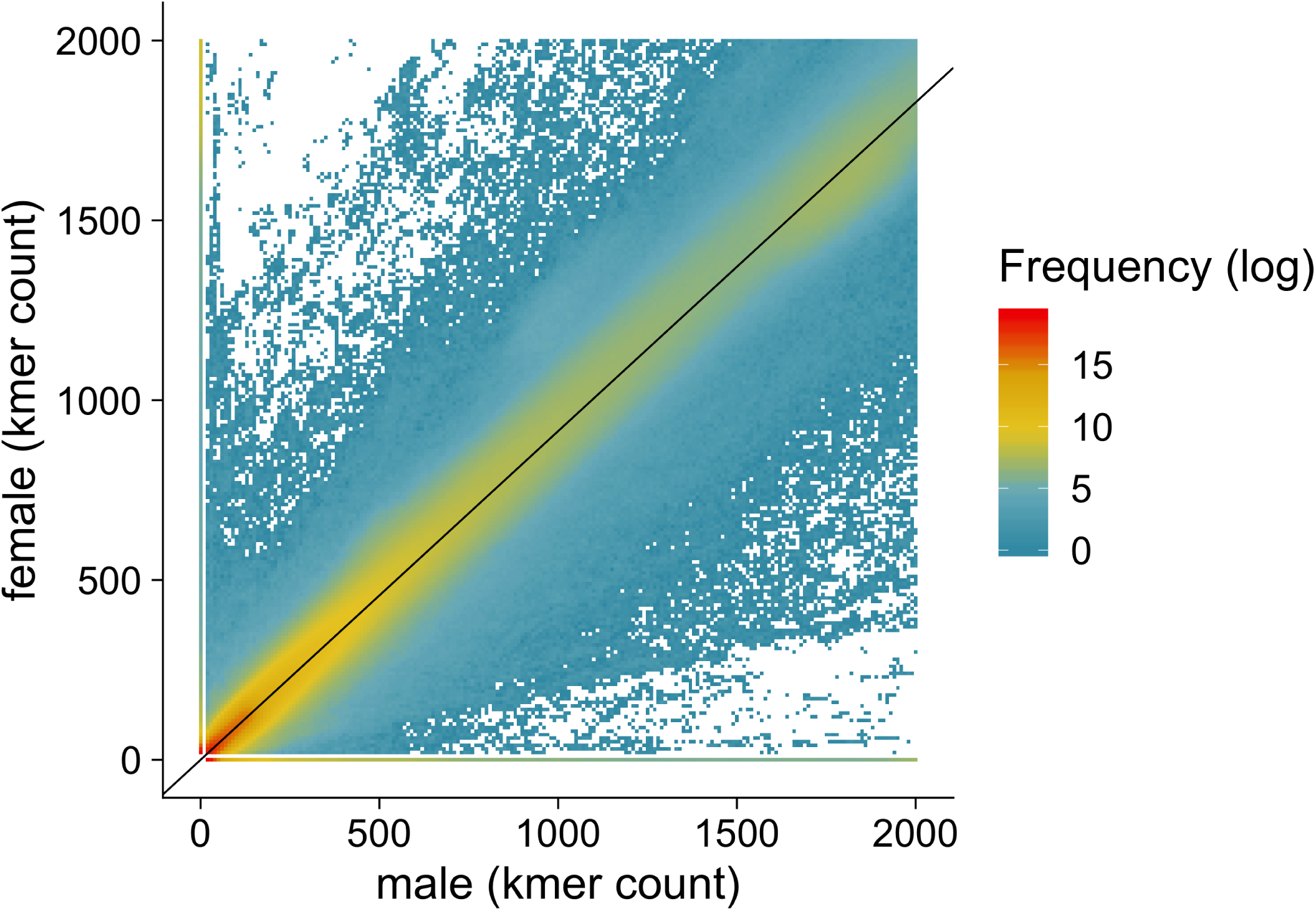
Comparison of the abundances of k-mers from the male and female sequence read pools of the Californian *A. palmeri* population. The color scale indicates the number distinct k-mers represented by one dot. The black line displays the null expectation for k-mer frequency in males and females, corrected for the number of individuals present in the male (n=35) and female (n=32) pool.

### Size estimation of the male specific region

We extracted all reads with identified male specific k-mers from the original reads data to assemble the male specific genome region. Only k-mers that were classified as male specific in both populations were used for the extraction of reads. The assembly process resulted in 5,774 contigs, which together consisted of 2,002,103 bp. The shortest contig was 151 bp long and the longest scaffold was 11,654 bp long (Fig.S2). After filtering out contigs with length below 152 (maximum length of a single read), we found 3,893 contigs comprising a total length of 1,748,724 bp. Based on the estimated genome size of 423 Mb (Stetter and Schmid 2017), the identified male specific region represents roughly 0.41% of the *A. palmeri* genome.

## Discussion

### Male specific genome region determines sex

The emergence of sex chromosomes is an evolutionary process that starts with the occurrence of single mutations leading to unisexual individuals and ultimately giving rise to highly heteromorphic chromosomes. Our results show that the genome of *A. palmeri* harbours a male specific region, genetically controlling the sex determination. The male specific region appears to not recombine with the analogous region on the X chromosome and has diverged sufficiently to not align to the hermaphrodite relative. The lack of alignment to the reference sequence consequently did not allow to detect allele frequency differences between pools. Although the male sequence is strongly diverged leading to recombination suppression between male and female chromosomes in the male specific region, an accumulation of repetitive sequences seems to not have occurred, as it would have led to an increase in chromosome size that could be cytologically identified. Such a cytological difference has not been observed in *A. palmeri* (Grant 1959).

To further characterize the evolutionary state of sex determination in the species, we estimated the size of the male specific region. Our assembly of the region suggests approximately 2 Mb. This size likely underestimates the true size of the male specific region. Our filtering removes highly redundant k-mers, as created by transposable elements (Michael 2014). Repetitive sequence is also present in other parts of the genome and would therefore not be classified as males specific. The high number of contigs potentially results from gaps due to non-specificity of the sequence, however, the stringent filtering across two independent populations revealed high confidence sequence specific to male individuals. Fine mapping of the male specific region and identification of encoded genes will deepen our knowledge about plant sex chromosomes. In contrast to the male specific k-mers, almost all female specific k-mers found in the California sample were removed after filtering for overlaps with the Kansas population. This confirms an XY system for *A. palmeri* as found by male specific k-mers. However, a very low number of apparently female specific k-mers persisted after filtering. These might be involved in female reproductive functions and could be further analyzed in future molecular studies.

### Evolution of dioecy in A. palmeri

The different stages of sex chromosome evolution may give us insight into the question as to when dioecy has evolved in a species. For example in the wild strawberry *Fragaria virginiana* male individuals coexist with females and hermaphrodites. Full dioecy is not yet evolved and the sex determining regions recombine to create unisexuals, hermaphrodites and even neuters (Spigler *et al*. 2008). This early stage of sex chromosome evolution indicates that the separation of sexes is rather young, whereas, the sex chromosomes in mammals and birds are very old (Cortez *et al.* 2014; Zhou *et al.* 2014). The Y (mammals) and W (birds) chromosomes of the heterogametic individuals of these taxa have undergone gene loss processes and finally became smaller than their X and Z chromosomes, respectively. Most heteromorphic sex chromosomes in flowering plants are increased in size as they accumulated transposable elements and other repetitive sequences. It is assumed, that this stage precedes gene loss (Ming *et al.* 2011). Our results show that the male specific region in *A. palmeri* is strongly diverged from the X region whilst cytological studies have not found a dimorphism between X and Y (Grant 1959). Therefore, we conclude that *A. palmeri* is at an early or intermediate stage of sex chromosome evolution.

The presence of cytologically homomorphic sex chromosomes that are nonetheless heteromorphic on the molecular scale has been shown in several other plant species (Liu *et al.* 2004; Khattak *et al*. 2006; Telgmann-Rauber *et al.* 2007), such as *Spinacia oleracea*, belonging to the same family as *A. palmeri*, i.e., Amaranthaceae.

Within the genus *Amaranthus, A. palmeri* and its dioecious sister species *A. arenicola* are placed in a monophyletic group (Stetter and Schmid 2017), in which all other species show monoecy (Wulff 1988; Costea *et al.* 2001; Asha *et al.* 2016). This suggests that dioecy has evolved independently within this group and is unlikely to be linked to the dioecious systems present in other *Amaranthus* species. To confirm this, we extracted reads with our set of male specific k-mers from whole genome sequencing data of *A. tuberculatus* individuals, another dioecious *Amaranthus* species, and compared the number of extracted reads (Fig. S3). We found no significant difference between male and female individuals, suggesting an independent evolution of dioecy in *A. palmeri*. This result agrees with the phylogentic distance between *A. palmeri* and *A. tuberculatus* which both cluster with several hermaphrodite species in genome-wide phylogenetic analyses (Stetter and Schmid 2017). Based on the phylogenetic relationship the sexual dimorphism in *A. tuberculatus* (and *A. australis* and *A. floridanus*) is older than dioecy in *A. palmeri*.

### Environmentally controlled sex determination

A recent study by Mesgaran *et al*. (2019) showed that the sex ratio in *A. palmeri* plants can be changed in response to water stress. Under well-watered conditions, males and females were present in a balanced ratio, while water stress resulted in an increased female to male ratio. These findings suggest that in *A. palmeri* sex is not purely genetically controlled, but behaves plastic in response to environmental factors. The increase in number of females in the water stress study suggests that genetic males can change their sex expression under altered environmental conditions. Hence, sex determining loci, found in the male specific regions and their X chromosome counterpart potentially interact with autosomal genes to integrate environmental factors and alter sex determination under stress conditions.

Our work constitutes the first step in understanding the underlying processes of sex determination in *A. palmeri*, but the exact mechanisms and the interaction with environmental factors remain to be discovered. The male specific k-mers identified here allow to diagnose the sex of individuals in early developmental stages. This permits future studies of environmental dependent sex alteration, as it will allow to distinguish between “genetic” sex and expressed sex of an individual plant. Furthermore, the 2 Mb male specific region can be used to fully assemble the *A. palmeri* Y chromosome. The different age and independence of dioecy evolution within the *Amaranthus* genus, make them a compelling system to understand the emergence of sexual dimorphism in plants. Further understanding of the sex determination in this invasive weed, can help develop novel ecological methods (e.g. shifting sex) to disrupt seed production in *A. palmeri* and hence reduce the impact of this invasive weed in cropping system.

## Acknowledgments

We thank Julia Kreiner for sharing records of sex expression of sequenced *A. tuberculatus* samples. MBM acknowledges the support from UC Davis New Research Initiatives and Interdisciplinary Research Grant and USDA-NIFA Hatch Funding (Project No. CA-D-PLS-2511-H). We acknowledge the support of the Deutsche Forschungsgemeinschaft under Germany’s Excellence Strategy – EXC-2048/1 – Project ID 390686111 to MGS.

## Author contribution

MBM and MGS designed and planned the study, MM and MBM provided plant material, AL performed molecular work, CJN and MGS analyzed the data. MGS supervised the study, and MT and MGS wrote the manuscript. All authors read and approved the manuscript.

## Data Availability

Sequencing data is available from the European Nucleotide Archive (ENA) under the project numbers PRJEB38372. Scripts used for the analysis are available through figshare 10.6084/m9.figshare.12326306

## Supplement

**Table S1.** Data Summary. ^1^ Paired end 2x 150bp reads. ^2^ Trimmed and quality filtered correctly paired reads. ^3^ 35-mers from filtered reads with 15 ≤ count ≤ 2000. ^4^ unique k-mers *times* number of each k-mer after correction with Kansas population. Values > 0 in opposite pools result from filter steps. For instance, more than 2,000 or below 20 k-mers in opposite pool

|  | AMA1 | AMA2 | AMA5 | AMA6 |
| --- | --- | --- | --- | --- |
| Population | California | California | Kansas | Kansas |
| Sex | male | female | male | female |
| Individuals | 35 | 32 | 25 | 35 |
| Raw read pairs<sup>1</sup> | 467,707,106 | 426,450,124 | 322,106,487 | 371,673,480 |
| Per pool coverage (X) | 331.71 | 302.45 | 228.44 | 263.60 |
| Per individual coverage (mean X) | 9.48 | 9.45 | 9.14 | 7.53 |
| Mapped to *A. hypochondriacus* (%) | 90.54% | 92.55% | 92.12% | 92.06% |
| Filtered paired reads<sup>2</sup> | 460,522,536 | 420,526,544 | 316,496,725 | 365,964,343 |
| Total k-mers<sup>3</sup> | 1,329,452,869 | 1,235,093,537 | 961,598,655 | 1,066,417,174 |
| k-mers/read pair (mean) | 2.89 | 2.94 | 3.04 | 2.91 |
| Sum male specific k-mers<sup>4</sup> | 17,259,830 | 0 | 13,168,285 | 52,698 |
| Sum female specific k-mers<sup>4</sup> | 83,397 | 324,143 | 12,994 | 249,012 |

**Figure S1.**
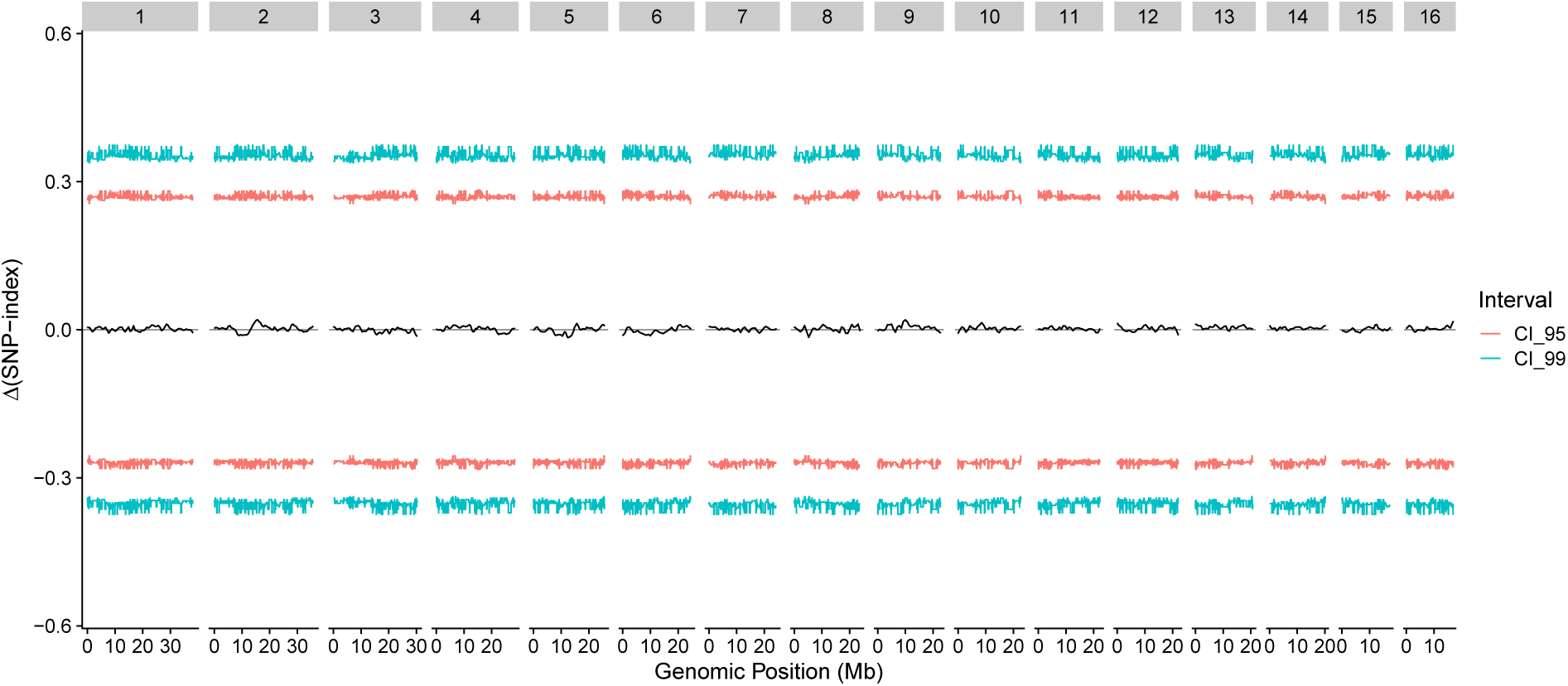
Differences in allele frequencies between male and female pools of the Kansas population along the genome, relative to the *A. hypochondiracus* reference. Red and blue lines represent 95% and 99% confidence intervals for frequency outliers.

**Figure S2.**
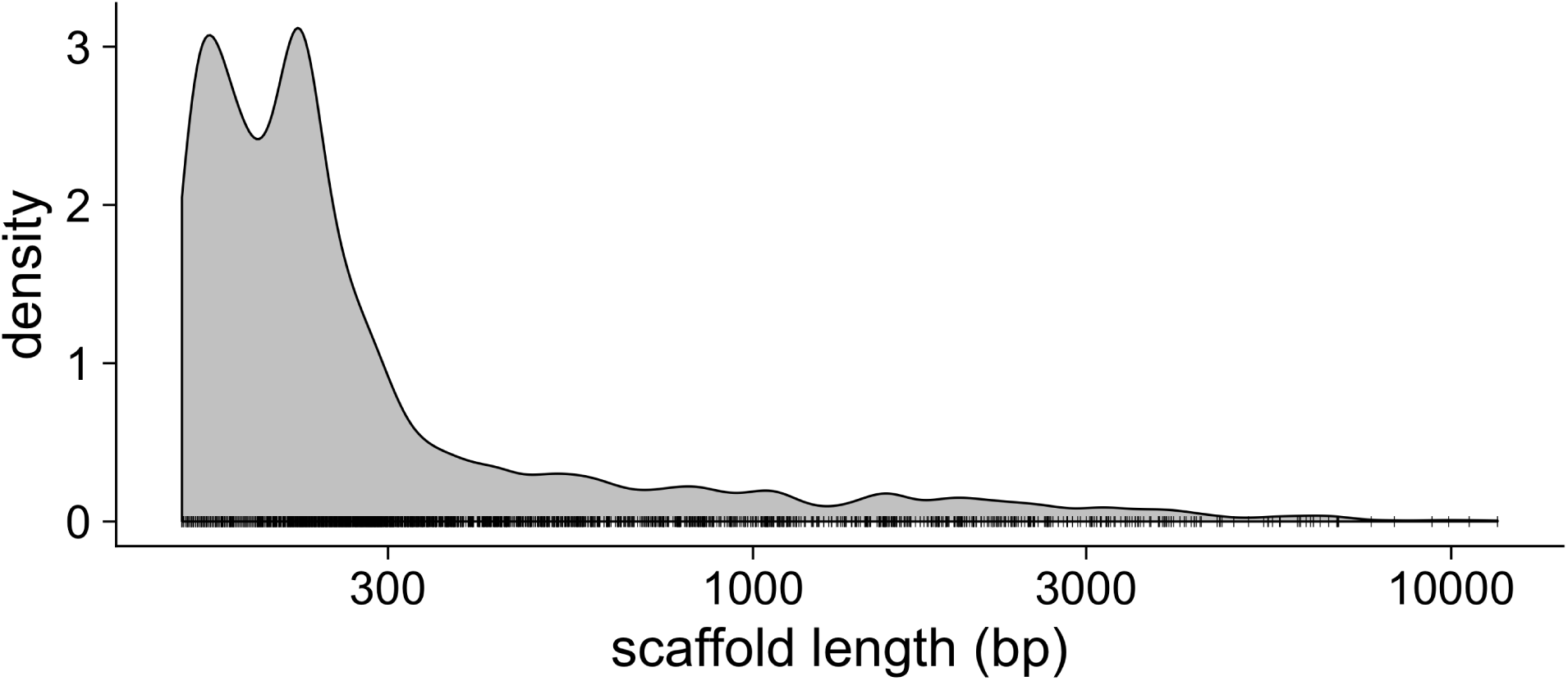
Scaffold size distribution. Size distribution of scaffolds assembled from sex specific reads. Ticks on the x-axis show value distribution

**Figure S3.**
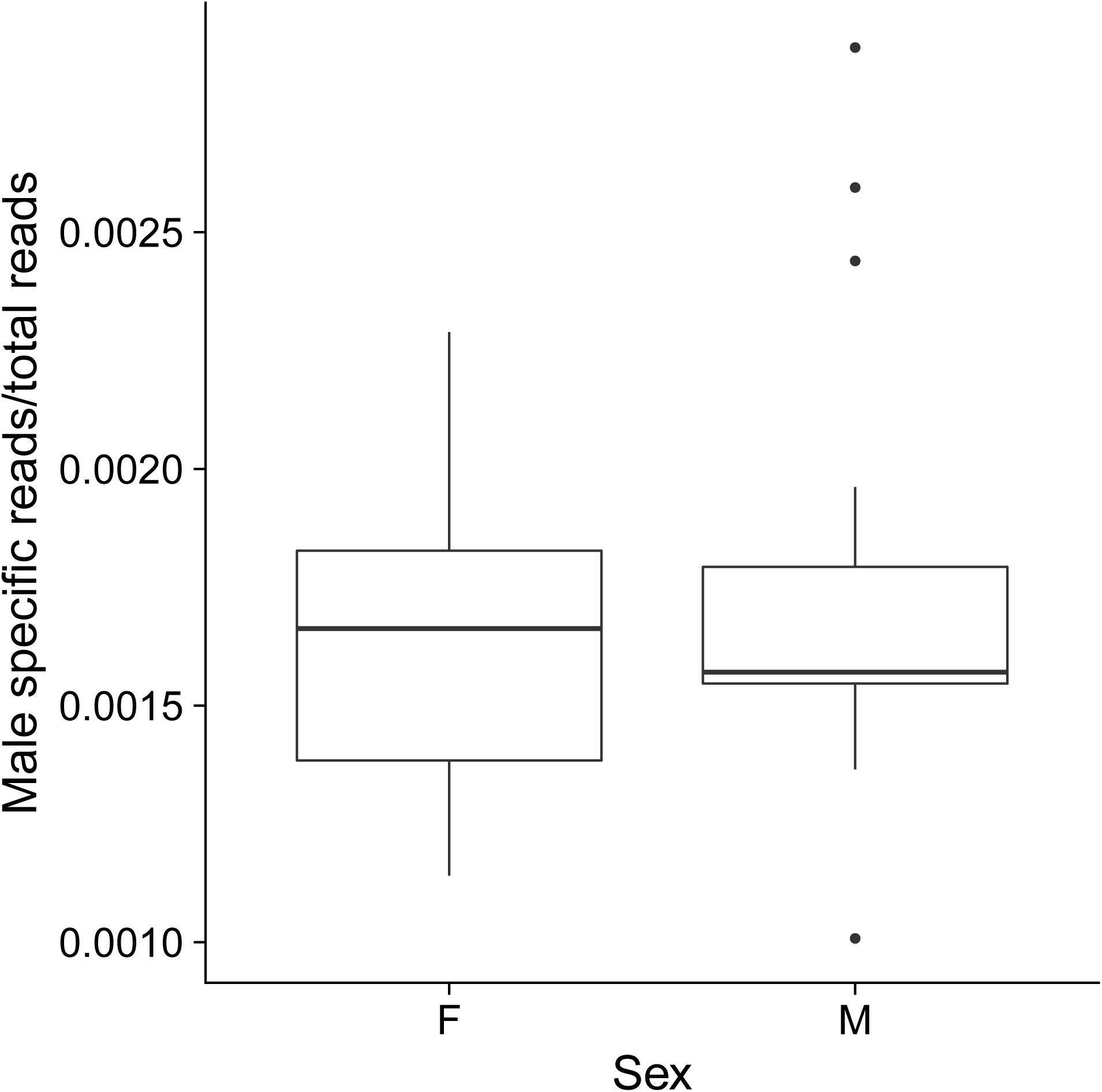
*A. tuberculatus* reads extracted. Ratio of *A. palmeri* male specific k-mer containing reads in *A. tuberculatus* samples. No significant difference between female (n=19) and male (n=25)

